# The interaction of XPG with TFIIH through p62 and XPD is required for the completion of nucleotide excision repair

**DOI:** 10.1101/2025.05.20.655039

**Authors:** Mihyun Kim, Eunwoo Jeong, Jiyoung Park, Miaw-Sheue Tsai, Chunli Yan, Ivaylo Ivanov, Hyun Suk Kim, Orlando D Schärer

**Affiliations:** Center for Genomic Integrity, Institute for Basic Science, Ulsan, 44919 Republic of Korea; Department of Biological Sciences, College of Information-Bio Convergence Engineering, Ulsan National Institute of Science and Technology, Ulsan, 44919, Korea; Biological Systems and Engineering Division, Lawrence Berkeley National Laboratory, Berkeley, CA, USA; Department of Chemistry, Georgia State University, Atlanta, GA, USA; Graduate School of Health Science and Technology, College of Information-Bio Convergence Engineering, Ulsan National Institute of Science and Technology, Ulsan, 44919, Korea; UPMC Hillman Cancer Center & Department of Pharmacology and Chemical Biology, University of Pittsburgh School of Medicine, Pittsburgh, PA, USA

**Keywords:** Nucleotide Excision Repair, XPG, TFIIH, Protein-Protein Interactions

## Abstract

Nucleotide excision repair (NER) is the key pathway for the removal of DNA damage induced by UV irradiation and chemotherapeutic reagents. Protein-protein interactions are crucial for the dynamic and coordinated assembly of the proteins involved at DNA lesion. Here we focus on the role of interactions between the multi-subunit helicase/translocase complex TFIIH and the 3’ endonuclease XPG. We show that XPG interacts with the p62 and XPD subunits of TFIIH through its long spacer region located in the middle of its split active site. We show the interactions between three acidic regions of XPG with the pleckstrin homology (PH) domain of p62 are of moderate importance for NER, while interactions with XPD are critical for activity. Combining mutations in the p62 and XPD interaction domains leads to additive defects in NER activity. Unexpectedly, we show that these interactions are not required for the recruitment of XPG, but rather for the formation of a catalytically competent NER complex and for triggering the incision 5’ to the lesion by ERCC1-XPF. Our studies provide fundamental insights into how interactions between TFIIH and XPG contribute to the NER pathway and more generally how modular protein-protein interactions control each step along the NER reaction coordinate.

## Introduction

Nucleotide excision repair (NER) is a multistep pathway involving more than 30 proteins that removes bulky lesions induced by UV light, environmental mutagens and chemotherapeutic agents from DNA (1,2). In global-genome (GG-) NER, XPC-RAD23B-CENT2 initiates NER by recognizing lesion-induced destabilization of duplex DNA and recruiting the TFIIH transcription/repair factor (3,4). In transcription-coupled (TC-) NER, a stalled RNA polymerase II (RNAPII) recognizes the damage. With help of CSA, CSB, UVSSA, ELOF1, and STK19, RNAPII is dislodged and TFIIH joins the complex (5–13). In both NER pathways, the core complex of TFIIH (XPB, XPD, p62, p52, p44, p34, and p8) opens the DNA using the XPB translocase and verifies the damage when the XPD helicase stalls at the lesion (14–19). XPA, RPA, and XPG are then recruited to stabilize the open DNA (3,15,20). Finally, ERCC1-XPF joins the NER complex and incises the DNA 5’ to the lesion, initiating repair synthesis and gap filling, 3’ incision by XPG, and sealing of the nick (21–24).

Protein-protein interactions among NER factors are essential for the assembly of the NER machinery and progression through the NER pathway (25). Several of these interactions, such as the ones between XPC and TFIIH (26–28), TFIIH and XPA (29,30), XPA and RPA (31–33), XPA and ERCC1 (34–36) have been characterized in some detail at the structural and functional levels. By contrast, less is known about the interaction domains that direct the XPG endonuclease to make the 3’ incision. Cellular data suggest that XPG joins the NER preincision complex following TFIIH and independently of and simultaneously with XPA and RPA (37,38). TFIIH and XPG have been shown to influence each other’s helicase and nuclease activity, respectively (30,39). Furthermore, it has been shown that XPG and XPC do not simultaneously co-exist in the NER preincision complex (15,20), suggesting that a handover from XPC to XPG is critical to the transition from the recognition to the preincision complex.

XPG is a member of the FEN-1 (flap structure specific endonuclease 1) family of nucleases with a split active site consisting of the highly conserved N-terminal (N)- and internal (I)-nuclease regions separated by a large spacer region of 680 amino acids (40). The spacer region in XPG has been shown to interact with TFIIH (41,42), but the information on specific interaction domains and their contribution to NER are limited.

Immunoprecipitation analysis showed that XPG binds to Pleckstrin homology (PH) of p62 (43). Additional insights into the interaction of XPG with p62 came from studies in yeast, where Rad2 (the *S. cerevisiae* XPG homolog) interacts with Tfb1 (the *S. cerevisiae* homolog of p62) via two acidic regions of Rad2 (residues 359-383 and 642-690) (44,45). Remarkably, this binding mode is similar to the interaction of human XPC with p62 and their *S. cerevisiae* homologs Rad4 with Tfb1, respectively. NMR titrations show that Rad2 can outcompete Rad4 for the Tfb1 binding as both Rad4 and Rad2 bind to the PH domain of Tfb1 via an acidic domain site (44,46,47).

Cross-linking mass spectrometry, though with limited experimental information, showed an interaction interface of the spacer region of XPG with the FeS cluster and Arch domains of XPD (30). Interactions between XPG and XPD were predicted in more detail in a computational model of the NER preincision complex (39). This model revealed a prominent interaction between two coiled-coil helices of XPG that emerge from the nuclease domain and mark the N- and C-terminal ends of the spacer region with the Arch and FeS domains of XPD.

Here, we combined the available information on the XPG-TFIIH interaction with sequence alignments and structure prediction to define interaction domains of XPG with the p62 and XPD subunits of TFIIH. Cellular and biochemical assays show that both interactions contribute to NER with a particularly prominent role for the XPG-XPD interaction. Unexpectedly, the interaction domains we characterize are not required for the association of XPG with the NER preincision complex, but instead are involved in the final assembly of the NER complex and licensing of the 5’ incision by ERCC1-XPF.

## Materials and Methods

### Plasmids and primers

HA tagged XPG cDNA in pWPXL and pFastBac vectors were used as starting constructs (22). Mutated pWPXL-XPG and pFastBac-XPG expression vectors were generated by site-directed mutagenesis from pWPXL-XPG WT and pFastBac-XPG WT using the KOD-mutagenesis kit (Toyobo, TYB-SMK-101). The following primers were used for inverse PCR mutagenesis to induce deletions or point mutations (altered codons underlined):

XPG Δ150-160 F: GATGAAAAAGAATGGCAAGAAAGAATGAATCA
XPG Δ150-160 R: TAAAGGAGGCAAAACATAGAGGTCGTTTTCTC
XPG Δ357-368 F: GCCCGTGGGAGGAACGCACCTG
XPG Δ357-368 R: GCTACTTCCCAGCAGGGCAGCTT
XPG Δ654-657 F: CAAAGTGTGATTAGTGATGAGGAACTT
XPG Δ654-657 R: ACTTCCATCAGATTCACTTTCTTCCG
XPG K115E F: TTTTTG<u>GAA</u>AGACAAGCCATCAAAACTGCC
XPG K108E/K111E R: TGT<u>TTC</u>CAGAAG<u>CTC</u>CTCTGTCGTTTTCCT
XPG F113A/L114A F: ACA<u>GCAGCA</u>AAAAGACAAGCCATCAAAACT
XPG L109A/L110A R: TTT<u>TGCTGC</u>CTTCTCTGTCGTTTTCCTG
XPG R124E F: <u>GAA</u>AGCAAAAGAGATGAAGCACTACCCA
XPG R124E R: GAAGGCAGTTTTGATGGCTTGTCTTTTCA
XPG R138E F: <u>GAA</u>AGAGAAAACGACCTCTATGTTTTGCC
XPG R138E R: AACTTGGGTAAGACTGGGTAGTGCTTC
XPG R168E F: <u>GAA</u>ATGAATCAAAAACAAGCATTACAGGAAG
XPG R168E R: TTCTTGCCATTCTTTTTCATCTTCCTCTTC

### Antibodies

The following antibodies were used for Western blots: anti-XPG (Bethyl, A301-484A), anti-HA (Abcam, ab9110), anti-Ku80 (Cell Signaling, 2753s), anti-XPD (Abcam, ab54676), anti-p62 (Santa Cruz, sc-48431), or anti-α-tubulin (Sigma Aldrich, T9026), goat anti-rabbit IgG antibody (Enzo, SAB-300-J), and goat anti-mouse IgG antibody (Enzo, SAB-100-J).

The following antibodies were used for local UV irradiation assays and slot-blot assays: anti-CPD (Cosmo Bio, CAC-NM-DND-001), anti-(6–4) PPs (Cosmo Bio, CAC-NM-DND-002), anti-HA (Abcam, ab9110), anti-XPF (Novus Biologicals, NBP2-58407), Goat anti-rabbit IgG Alexa Fluor^TM^ 488 (Thermo Fisher Scientific, A11008), and Cy™3 AffiniPure Goat Anti-Mouse IgG (Jackson Immunoresearch, 115-165-146).

### XPG protein expression and purification

His-tagged XPG WT and mutant proteins were expressed in Sf9 insect cells (0.4 L culture) as described previously (48). The harvested cells were resuspended in 40 mL of Lysis buffer (0.01 M PBS, pH7.4, 500 mM NaCl, 1 mM PMSF, 0.5% NP-40, PI tablet, and 1 mM 2-mercaptoethanol). After incubation on ice for 20 min, the lysate was homogenized using a Dounce homogenizer and clarified by ultracentrifugation (40,000 g for 45 min). The supernatant was incubated with nickel-nitrilotriacetic acid-agarose beads (Ni-resin) (Qiagen) in the presence of 10 mM imidazole for 1 h. The beads were collected by low-speed centrifugation and resuspend in 10 mL of washing buffer (10 mM PBS at pH7.4, 500 mM NaCl, and 0.1% NP-40) containing 10 mM imidazole and packed in column. The column was washed stepwise with 10 mL of washing buffer containing 20 mM imidazole followed by 10 ml of 40 mM imidazole. XPG was eluted with Lysis buffer containing 250 mM imidazole. The eluent was dialyzed overnight against dialysis buffer (20 mM Tris-HCl at pH 7.5, 100 mM NaCl, 5% glycerol, 0.1% NP-40, 1 mM 2-mercaptoethanol) at 4°C and the dialyzed protein applied to a SP Sepharose column (Amersham Biosciences). The column was washed two times with 10 column volume (CV) of SP buffer (20 mM Tris-HCl at pH 7.5, 100 mM NaCl, 5% glycerol, 0.1% NP-40) containing 100 mM NaCl and 150 mM NaCl, respectively. The proteins were eluted 6 times with 0.5 CV of SP buffer containing 800 mM NaCl. The eluted proteins were further purified through gel filtration (Superose 6 Increase 10/300 GL, Amersham Biosciences) pre-equilibrated in GF buffer (100 mM Tris-HCl at pH 7.5, 300 mM KCl, 0.1% NP-40, 1% glycerol). The XPG containing fractions were pooled and glycerol was added to a final concentration of 10%. Protein concentration was measured using Bradford assay and stored at −80°C. Around 350 µg of protein was obtained from 400 mL culture.

### p62-PH protein expression and purification

Gene of the PH domain of p62 (residues 1-108) was synthesized by ‘Bionics’ and the sequence was inserted to pBG100 vector to generate pBG100 p62-PH plasmid. The plasmid was transformed into *Escherichia coli* BL21/(DE3) Rosetta pLysS and cells were grown in LB medium containing 50 µg/ml kanamycin to OD_600_ = 0.6 at 37°C, then to OD_600_ = 1.2 at 18°C, and p62-PH expression induced with 1 mM isopropyl β-D-thiogalactoside (IPTG) at 18°C overnight. Cells were collected by centrifugation at 6,500 rpm for 20 min and all subsequent purification steps were performed at 4°C. The cell pellet from 2 L of culture was resuspended in 30 mL of lysis buffer (100 mM Tris-HCl at pH 8.0, 500 mM NaCl, 20 mM imidazole, 5 mM 2-mercaptoethanol, 200 µg/ml lysozyme, 0.5 mM PMSF, 1 mM benzamidine, and 10% glycerol, and protease inhibitor (Roche, 11697498001)) and lysed with 20 strokes in a Dounce homogenizer and sonicated with an amplitude of 30 (5 sec on/10 sec off) for 2 min. The lysate was clarified by centrifugation at 20,000 rpm for 40 min and filtration of the supernatant through a 0.45 µm PVDF syringe filter 3 times. The supernatant was incubated with 2 mL of Ni-resin pre-equilibrated with Ni-loading buffer (100 mM Tris-HCl at pH 8.0, 500 mM NaCl, 20 mM imidazole, 5 mM 2-mercaptoethanol, and 10% glycerol) for 60 min at 4°C. The resin was washed twice with Ni-wash buffer (100 mM Tris-HCl at pH 8.0, 500 mM NaCl, 30 mM imidazole, 5 mM 2-mercaptoethanol, and 10% glycerol) and eluted with Ni-elution buffer (100 mM Tris-HCl at pH 8.0, 500 mM NaCl, 250 mM imidazole, 5 mM 2-mercaptoethanol, and 10% glycerol). The proteins were further applied to a HiLoad 16/600 Superdex 75 pg column and eluted with 50 mM Tris-HCl at pH 8.0, 150 mM NaCl, 1 mM dithiothreitol, and 10% glycerol. p62-PH eluted at 70-80 mL and was obtained at a concentration of ∼9.5 mg/mL, yielding a total of ∼5 mg per liter of cell culture.

### Fluorescence anisotropy

The following FITC-labeled peptides were obtained from Cusabio with a declared purity of <95%.

XPC 124-142: EEEEESENDWEEVEELSEP (XPC peptide sequence from Okuda et al, 2015)
XPG 150-168: QEEEKHSSEEEDEKEWQER
XPG 355-373: SSSEEELESENRRQARGRN
XPG 642-665: EIDSEESESDGSFIEVQSVISDEE

Each peptide (0.5 mg) was dissolved in 20 µL of DMSO and diluted to 0.5 mM with buffer F (20 mM Tris-HCl at pH 8.0 and 20 mM NaCl). FITC-labeled peptide (250 nM) was incubated with indicated concentrations of His-p62-PH protein for 2 min. After 2 min the fluorescence intensity was measured at 490 nm excitation wavelength on a fluorescence spectrophotometer (Cary Eclipse). The unbound fraction was calculated using the following equation.

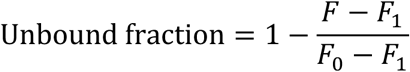

(F_0_: Fluorescence intensity of 0 nM of p62-PH, F_1_: Fluorescence intensity of 1000 nM of p62-PH)

The data was analyzed by ‘Graphpad Prism 5’ to determine the Kd value using saturation curve.

### Immunoprecipitation

Cells were lysed with buffer X (100 mM Tris-HCl at pH 8.5, 250 mM NaCl, 1 mM EDTA, and 1% NP-40, 5 mM MgCl_2_) supplemented with phosphatase inhibitor PhosSTOP (Roche), complete protease inhibitor cocktail (Roche), and 50 units of Benzonase for 1 h at 4°C while rotating. Lysates were cleared by centrifugation at 15,000 rpm for 25 min at 4°C. HA-tagged XPG proteins were incubated with anti-HA agarose affinity beads (Sigma) for 1 h at 4°C with rotation. The beads were washed three times using buffer X, and the bead-bound proteins were eluted by boiling at 95°C for 5 min. Proteins eluted from the beads were loaded onto SDS-PAGE and analyzed by immunoblotting. Samples were separated by electrophoresis on a 4-12 % Bis-Tris gels (Thermo Fisher Scientific) at 80 V for 95 min, transferred to PVDF membrane (Amersham Hybond 0.2 µm PVDF) at 300 mA for 80 min using the mini-protean tetra system (Bio-Rad) at 4°C, and blocked with 5% skim milk in Tris-buffered saline containing 0.1% Tween-20 (TBST) for 1 h at room temperature (RT). Rabbit polyclonal XPG antibodies (1:1,000), Mouse monoclonal XPD antibodies (1:3,000), p62 antibodies (1:1,000), or α-tubulin (1:5,000) were added to the TBST and incubated overnight at 4°C. The blot was then incubated with goat anti-rabbit IgG antibody (1:3,000) or goat anti-mouse IgG antibody (1:1,000) for 1 h at RT and the bands visualized using a chemiluminescent substrate (Thermo Fisher Scientific, 34577) and ChemiDoc touch imaging system (Bio-Rad, 1708370).

### AlphaFold 2 prediction of XPG-p62

Alphafold (v2.3.2) was installed on a local Ubuntu (v24.04 LTS) environment equipped with RTX 4090 24Gb and RAM 64Gb (49). The amino acid sequence of XPG (UniProtKB: P28715) and p62 (UniProtKB: P32780, A.A 1-108) in FASTA format were used as input. The parameters were set as default except max_template_date=1900-04-01. We used the top-ranked model with the highest confidence scores calculated by Alphafold and analyzed with PyMol (Schrödinger Inc).

### Generation of cell lines expressing mutant XPG proteins

SV40-transformed human fibroblasts XP3BRSV (XPG-deficient cells) were cultured in Dulbecco’s Modified Eagle’s Medium (DMEM, Cytiva SH30243.01) supplemented with 10% fetal bovine serum (FBS, Millipore TMS—13-BKR) and penicillin streptomycin (P/S, Gibco) at 37°C in the presence of 5% CO2. For lentiviral production, 0.75 µg of pWPXL expression vector containing XPG cDNA, 2.25 µg of pMD2.G envelop plasmid, and 2.25 µg of psPAX2 packaging plasmid were transfected into 293T cells using Lipofectamine 3000 (Thermo Fisher Scientific, L3000001) according to established protocols and lentivirus was harvested 1 day post transfection (50). One day before transduction,XP3BRSV cells were seeded in 6-well plates at 50% confluency and incubated with lentivirus with a multiplicity of infection (MOI) of 2 for 24 h. Then, 2000 cells were seeded on the 10 cm plate to screen for single colonies for 7 days and clones expressing comparable XPG level were selected.

### Western blot

Cells were lysed with RIPA buffer (50 mM Tris-HCl at pH 8.0, 150 mM NaCl, 5 mM EDTA, 1% Triton X-100, 0.1% SDS, 0.05 g/mL sodium deoxycholate, 1 mM phenylmethylsulfonyl fluoride (PMSF), 5 mM MgCl_2_), supplemented with protease inhibitors (Roche). Protein concentration was measured using Bradford assays (Thermo Fisher Scientific). After addition of LDS sample buffer containing 2.5% of 2-mercaptoethanol (4x, Invitrogen), the samples were heated to 95°C for 5 min. Samples containing 25 µg of protein were separated by electrophoresis on 3-8% Tris-Acetate gels (Thermo Fisher Scientific). Gel running, transfer, and detection with antibodies were proceeded as above.

### Clonogenic cell survival assay

Native and transduced XP3BRSV cells were cultured as described above. 2000 cells were seeded in a 6-cm dish the day before UV irradiation. Cells were treated with 0, 1, 2, 4 J/m^2^ of UV-C, and grown for 10 days. Cells were fixed with 4% paraformaldehyde for 15 min and stained with 1.5% of methylene blue for 2 h. After washing with water, colonies containing more than 25 cells were counted. The survival rate was normalized to the number of colonies of non-UV treated cells. For quantification, more than 25 cells were considered as one colony. Each experiment was conducted as 3 independent replicates and error bar represents SEM of all experiments.

### Co-localization assay of XPG-CPD and XPF-CPD

5.0 × 10^3^ cells were seeded on slide cover glass (Neuvitro, GG-22-PDL) the day prior to UV irradiation. For UV irradiation, medium was removed and cells were washed with PBS. The cells were covered with 5 µm isopore membrane (Merck, TMTP04700) and irradiated with UV-C at a dose of 100 J/m^2^. Cells were then incubated in growth medium for 0.5-24 h. Cells were then lysed with cold hypotonic buffer (10 mM Tris-HCl at pH 8.0, 2.5 mM MgCl_2_, 10 mM β-glycerophosphate, 0.5% Igepal CA-630 (Sigma Aldrich, I8896), 0.2 mM PMSF, 0.1 mM Na_3_VO_4_) for 15 min on ice and washed with hypotonic buffer without Igepal for 4 min at RT. Cells were fixed in 2% formaldehyde in DPBS for 5 min at RT. After washing twice with 0.1% Triton X-100 in DPBS for 5min, cells were stored in 70% ethanol at −20°C. After removal of the ethanol, cells were washed with DNase I digestion buffer (10 mM Tris-HCl at pH 8.0, 5 mM MgCl_2_) for 2 min and then the DNA digested with 20 U of DNase I (cat #: D4527, Sigma-Aldrich) in 1 mL of digestion buffer for 40 sec at RT. EDTA (500 mM) was added to a final concentration at 10 mM EDTA to stop the reaction. The solution was removed and cells were washed twice with 10 mM EDTA in DPBS for 5 min. After treatment with blocking buffer (1% BSA, 0.2% Tween-20), cells were incubated with diluted primary antibodies, including CPD (1:1,500) and HA-tag (1:250) or XPF (1:100), in blocking buffer for 2 h. Following 3 washing steps with DPBS containing 0.2% Tween-20, cells were incubated with Goat Anti-Rabbit IgG Alexa Fluor^TM^ 488 (1:1,000) and then with Cy™3 AffiniPure Goat Anti-Mouse IgG (1:1,500) antibodies in blocking buffer and stained with Vectashield with DAPI (Vector laboratories, H-1200-10).

### Slot-blot assay

Cells were irradiated with 5 J/m^2^ of UV-C and collected after different repair time (0, 2, 4, and 8 h for (6–4) PPs and 0, 8, 24, and 48 h for CPDs). Genomic DNA was prepared from harvested cells using QIAamp DNA mini kit (Qiagen) according to the manufacturer’s protocol. After adjusting genomic DNA concentration, DNA was denatured with 7.8 mM EDTA (for (6–4) PPs) or with 0.4 M NaOH and 10 mM EDTA (for CPDs). The denatured DNA was boiled at 95°C for 10 min. DNA samples (200 ng of (6–4) PPs or 100 ng of CPDs) were neutralized by adding an equal volume of 2 M ammonium acetate (pH 7.0) and vacuum-transferred to a pre-washed nitrocellulose membrane using a BioDot SF microfiltration apparatus (Bio-Rad). Each well was washed 2 times with SSC buffer (0.15 M sodium chloride and 15 mM sodium citrate, pH 7.0). The membrane was removed from the apparatus, rinsed twice with SSC buffer, air-dried, baked under vacuum at 80°C for 2 h, and blocked with 5% skim milk in PBS. For lesion detection, the membrane was incubated with (6–4) PPs (1:2,000) or CPDs (1:3,000) antibodies at 4°C overnight and then incubated with anti-goat IgG mouse antibody (1:2,500 for (6–4) PPs or 1:5,000 for CPDs) for 1 h at RT. The blot was visualized with the ECL system (Thermo Fisher Scientific, 34577) and the total amount of DNA loaded on the membrane was visualized with SYBR-gold (Thermo Fisher Scientific, S11494) staining.

### CometCHIP assays

Native and transduced XP3BRSV cells in growth medium were seeded onto 1.5% agarose Comet-Chip (Sigma, A4018) and incubated with 4 mM HU and 40 µM AraC for 1 h prior to UV irradiation. Following 5 J/m^2^ UV-C irradiation, cells were further incubated with 4 mM HU and 40 µM AraC at 37°C for 0, 1, 2, and 4 h. Comet-Chips were then covered with 1% LM Agarose (Trevigen, 4250-050-02) and incubated with lysis buffer (Trevigen, 4250-050-01) overnight at 4°C. Following lysis, DNA was partially unwound in alkaline electrophoresis buffer (0.3 M NaOH and 1 mM EDTA, pH 13.5) for 40 min at 4^°^C, and subjected to electrophoresis for 30 min at 1 V/cm and 300 mA at 4°C. The chips were washed with 0.4 M Tris-HCl, pH 7.4 and stained with SYBR Gold (Thermo Fisher Scientific, S11494?) according to manufacturer’s protocol. Images were captured on a fluorescence microscope (Olympus, BX53), and comet tail lengths were analyzed using Comet analysis software (Trevigen, 42600-000-CS) (51).

### Nuclease activity assay

The fluorescently labeled 90 mer oligonucleotide, FAM-5 ′ - CCAGTGATCACATACGCTTT GCTATTCCGGTTTTTTTTTTTTTTTTTTTTTTTTTTTTTTCCGTGCCACGTTGTATG CCCACGTTGACCG-3′ was annealed with a 1.5-fold excess of the unlabeled oligonucleotides, 5’ - CGGTCAACGTGGGCATACAACGTGGCACGGTTTTTTTTTTTTTTTTTTTTTTTTTTTTTTCCGGAATAGCAAAGCGTATGTGATCACTGG-3’, in 10 mM Tris-HCl, pH 7.5 and 50 mM NaCl by heating for 5 min at 95°C and slowly cooling to RT. The resulting bubble DNA substrate was incubated with 5-40 nM XPG in cleavage buffer (25 mM Tris-HCl at pH 6.8, 10% glycerol, 0.5 mg/ml bovine serum albumin, 30 mM KCl, 2 mM MgCl_2_) at 30°C for 30 min. The reactions were stopped by the addition of formamide loading buffer and heating for 5 min at 95°C. The samples were analyzed by 15% denaturing PAGE, which was run at 15 mA for 1 h using 0.5X TBE buffer. The bands were visualized using Typhoon RGB imager.

### *In vitro* NER activity assay with purified proteins

A plasmid containing a GTG 1,3-Cisplatin lesion was incubated with purified NER proteins in the absence or presence of purified wild-type or mutant XPG proteins. For each reaction, 5 nM of XPC-RAD23B, 10 nM of TFIIH, 20 nM of XPA, 41.6 nM of RPA, 27 nM of XPG, and 13.3 nM of XPF-ERCC1 was used. All proteins were >95% pure and produced as previously described for XPC-RAD23B (52), TFIIH (53), RPA (54), XPG (48), and ERCC1-XPF (55). The reactions were conducted in repair buffer containing 45 mM HEPES-KOH at pH 7.8, 5 mM MgCl_2_, 0.3 mM EDTA, 40 mM phosphocreatine (di-Tris salt, Sigma), 2 mM ATP, 1 mM DTT, 2.5 μg BSA, 0.5 μg creatine phosphokinase (Sigma), and NaCl (to a final concentration of 70 mM). The purified NER proteins and repair buffer, total volume of 9 μl were pre-warmed at 30°C for 10 min. 1 μL of plasmid containing GTG 1,3-Cisplatin (25 ng/μl) was added to reaction mixture and incubated at 30 °C for different incubation times (0, 5, 10, 20, 45, and 90 min). 0.5 μL of 1 μM of a 3′-phophorylated oligonucleotide for product labeling was added and the mixture heated at 95°C for 5 min. The mixture was allowed to cool down to RT for 15 min, followed by adding 1.2 μL of a Sequenase/[α-^32^P]-dCTP mix (0.25 units of Sequenase and 2.5 μCi of [α-^32^P]-dCTP per reaction) and incubating at 37°C for 3 min. Then 1.2 μL of dNTP mix (100 μM of each dATP, dTTP, dGTP; 50 μM dCTP) was added and incubated for another 12 min. The reactions were stopped by adding 12 μL of loading dye (80% formamide/10 mM EDTA) and heating at 95°C for 5 min. Sample (7 μL) was loaded on 14% sequencing gel (7 M urea, 0.5x TBE) and ran at 45 W for 2.5 h. The reaction products were visualized using a PhosphorImager (Amersham Typhoon RGB, GE Healthcare Bio-Sciences). Two independent repetitions were performed. The NER products were quantified using ImageQuant TL and normalized to the amount of NER product formed with WT-XPG at 90 min.

## Results

### Predicted interfaces of p62-PH and XPD with XPG

The XPG endonuclease contains two highly conserved nuclease domains (**Figure 1A**) that are connected by a non-conserved and unstructured spacer region of ∼600 amino acids. This spacer region is known to mediate interactions with other proteins including TFIIH (40–42), but the details and functional significance of these interactions have so far remained elusive. We hypothesized that the interaction of XPG with two TFIIH subunits, p62 and XPD, may be important (**Figure 1B**). Studies in *S. cerevisiae* showed that the PH domain of Tfb1, the yeast homolog of p62, interacts with two acidic regions of Rad2 (**Figure 1C**) (44). We aligned eukaryotic XPG sequences and identified three domains rich in acidic amino acids: XPG_150-160_, XPG_357-368_, and XPG_654-657_ (**Figure 1A and S1A**). Each domain was highly conserved from xenopus to human. In particular, an FIEV motif (XPG_654-657_) shared similar sequences in Rad2 (FEDV) and in a p62-PH-interacting motif in UVSSA (FVEV motif) (28) (**Figure 1A and S1A**). We carried out an AlphaFold 2 modeling of the interaction of these three acidic regions with p62-PH, and the model predicted that XPG_654-657_ binding to p62-PH was similar to the Rad2-Tfb1 interaction, while XPG_150-160_- and XPG_357-368_-bound p62 appeared less specific (**Figure 1D**). To assess the importance of the interaction between XPG and p62-PH for NER, we generated XPG variants with deletion of the three individual acidic domains and with deletion of all three acidic domains. (**Table 1**).

**Figure 1.**
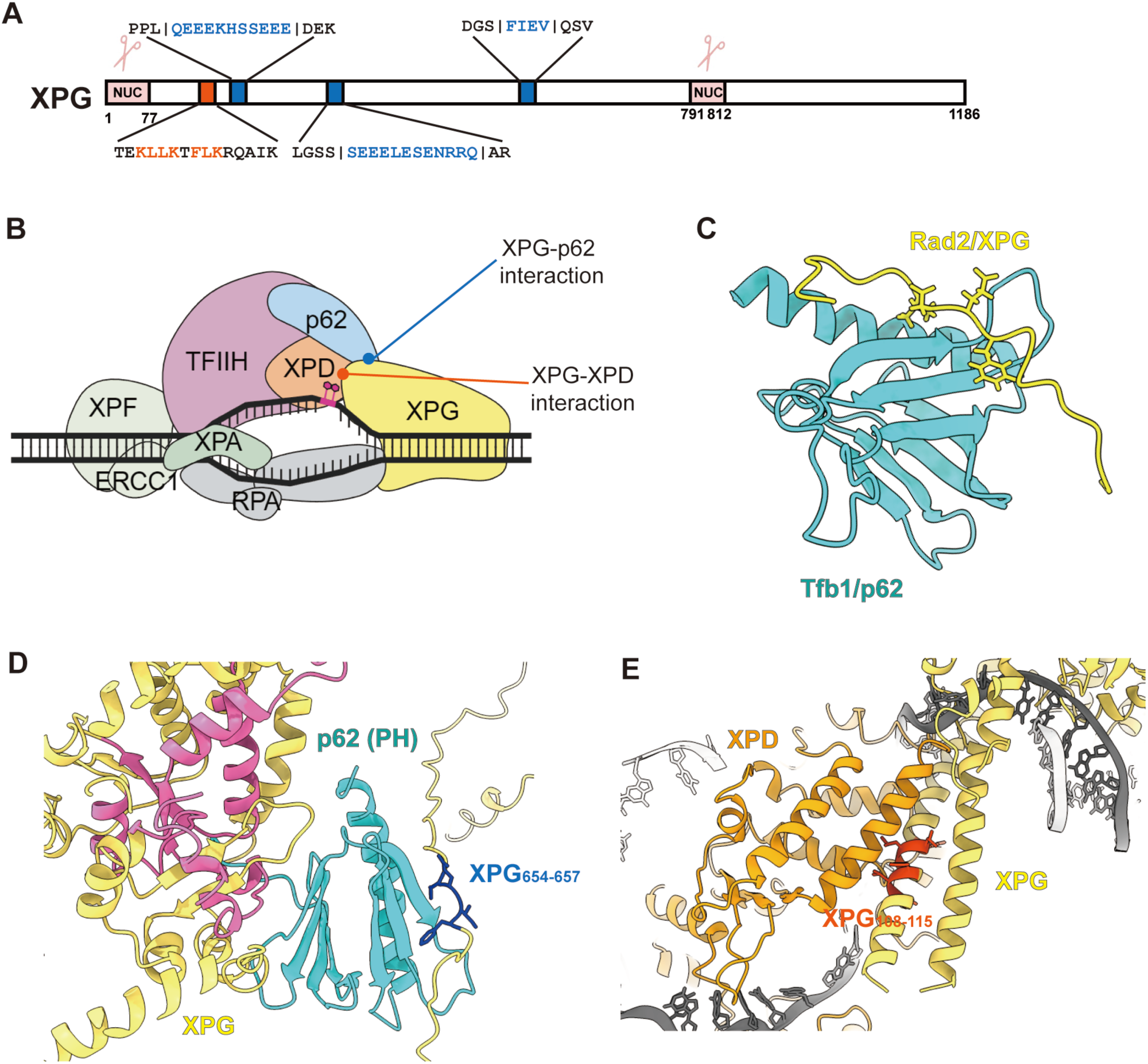
Interaction of XPG with p62 and XPD. (A) The domain map of the XPG protein with nuclease (NUC) domain and interaction domains with p62 (blue) and XPD (orange) of TFIIH highlighted. (B) Scheme of pre-incision complex. XPG interactions with p62 and XPD were indicated with blue and orange lines. (C) Structure of a Rad2 (XPG) peptide (yellow) bound to Tfb1-PH (p62) protein (cyan) from PDB: 2LOX. (D) AlphaFold 2 prediction of XPG and p62-PH. XPG is yellow with its nuclease domain highlighted pink; XPG_654-657_ is blue. Accuracy of prediction 45< pLDDT value of XPG_654-657_ <55. (E) Interaction of the gateway helical domain of XPG with XPD in the computational model of the preincision complex (Yu, 2024). DNA is grey, XPG yellow, and XPD orange. Residues XPG_108-115_ predicted to interact with XPD are highlighted in red-orange.

Based on recent computational modeling of the NER preincision complex, we predicted that a key interaction between XPD and XPG occurs between the Arch domain of XPD and the two long α-helices adjacent to the nuclease domain of XPG. (**Figure 1E**) (56). Within this helix, residues Lys108, Leu109, Leu110, Lys111, Phe113, Leu114, and Lys115 were predicted to bind XPD_257-271_ (**Figure 1E**) (56). Additionally, residues Arg124, Arg138, and Arg168 of XPG were reported to bind XPD based on crosslinking mass spectrometry analysis (30). To disrupt the interaction between XPG and XPD, a series of charge reversal mutations Lys/Arg to Glu or substitutions of Leu/Phe to Ala were introduced in pairs of 3-4 residues (**Table 1**). We used ddMut (57) to estimate the effect of these mutations on protein stability and found that these mutations do not affect overall protein stability (**Figure S1B**).

### Disruption of the interaction of XPG with p62-PH results in mild sensitivity to UV

To probe the interaction of XPG with p62-PH, we designed peptides containing the putative p62-PH interacting domain of XPG (**Figure 1A**) and measured their interaction with p62-PH using fluorescence anisotropy. We used a known p62-PH binding peptide of XPC as a positive control (47) (**Figure 2B**). FITC-labeled XPC and XPG peptides were incubated with the purified PH domain of p62 (residues 1-108) and the reduction of fluorescence intensity with increasing concentrations of p62-PH was measured (**Figure 2B**). A dissociation constant (Kd) of 98.7± 13.5 nM was determined for the XPC_124-142_ peptide, similar to the value previously measured by isothermal titration calorimetry (ITC) (57.5 ± 5.5 nM) (47) (**Figure 2B**). The XPG_150-168_ and XPG_355-373_ bound to p62-PH with comparable affinity as the XPC_124-142_ peptide (39.8 nM and 29.5 nM, respectively). Unexpectedly, the XPG_642-665_ peptide showed a higher Kd value (167.6 ± 66.1 nM) for p62-PH than XPG_150-168_ and XPG_355-373_, despite a good binding fit predicted by AlphaFold 2 (**Figure 1D and 2B**).

**Figure 2.**
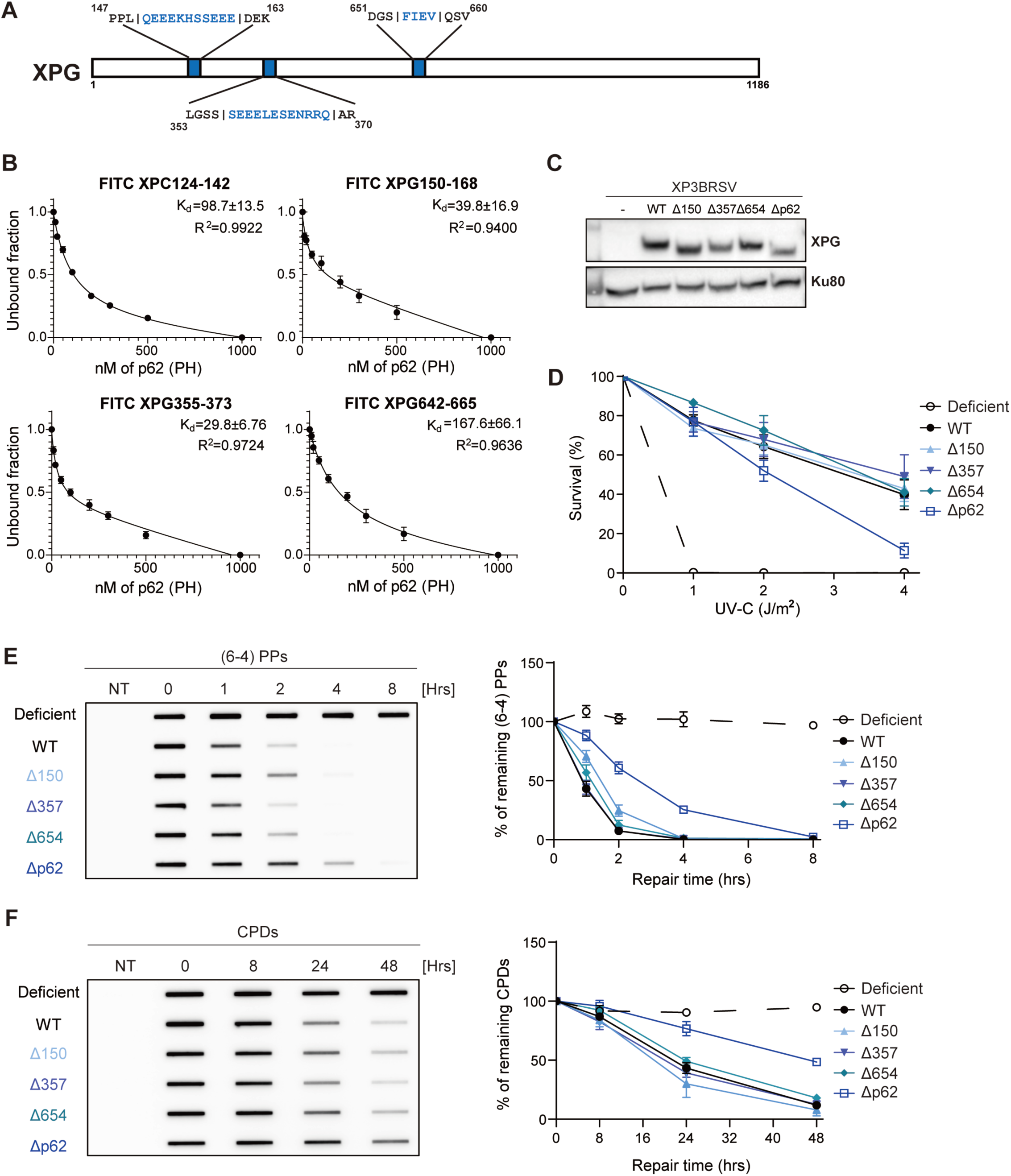
Mutations in the p62 interaction domains of XPG cause moderate defects for UV-induced DNA damage repair. (A) The XPG map with p62 interaction domains in blue. (B) Fluorescence anisotropy measurement of FITC-XPC and XPG peptides incubated with p62-PH protein. 250 nM of the peptide was incubated with the indicated amount of protein. The data represents the mean ± SEM of 4 independent experiments (N=2 with His-p62-PH, N=2 with p62-PH). (C) Western blot of XPG-Δ150, -Δ357, -Δ654, and -Δp62 cells generated by lentiviral transduction and clonal selection to reconstitute XPG variants in XP3BRSV (XP-G patient) cells. (D) Clonogenic survival assays. Cells were treated with UV-C (0, 1, 2, 4 J/m^2^), incubated for 10 days, stained with methylene blue, and colonies counted. Survival rates were normalized to the number of colonies of non-treated cells and shown as a mean ± SEM of 3 independent experiments. (E) Kinetics of (6–4) PPs repair determined by slot-blot assays. Cells were irradiated with UV-C (5 J/m^2^), genomic DNA isolated at the indicated time points, and the adduct levels determined using an anti-(6–4) PPs antibody. For quantification, band intensities were normalized to the WT band at 0 h and represented by a mean ± SEM of 3 independent experiments. (F) Kinetics of CPDs repair determined by slot-blot assays. Cells were irradiated with UV-C (5 J/m^2^), genomic DNA isolated at indicated time points, and the adduct levels determined using an anti-CPDs antibody. For quantification, band intensities were normalized to the WT band at 0 h and shown as a mean ± SEM of 3 independent experiments.

Next, we asked if the p62 interaction of XPG is required for the repair of UV-induced DNA damage. We generated cells expressing XPG with deletion mutations in the three individual (Δ150, Δ357, Δ654) or combined (Δp62) p62-interacting domains by lentiviral transduction of XP-G patient XP3BR cells and selected clones that expressed WT and mutant XPG proteins at comparable levels (**Figure 2A and 2C**). We first tested the sensitivity of these cells to UV-C irradiation. XPG-deficient cells were hypersensitive to UV-C irradiation, whereas more than 50% of the XPG-WT cells survived after exposure to 4 J/m^2^ of UV-C irradiation. Cells expressing XPG-Δ150, XPG-Δ357, or XPG-Δ654 showed similar sensitivity compared to XPG-WT (**Figure 2D**). Deletion of all three p62-interacting domains of XPG (XPG-Δp62) yielded cells with moderate sensitivity (∼3-fold) to UV (**Figure 2D**).

We then measured the repair kinetics of UV-induced damage using slot-blot assays. Cells were globally irradiated with UV-C (5 J/m^2^) and the amount of (6–4) PPs or CPDs was measured at different time points post UV irradiation (**Figure 2E, 2F, S2A, and S2B**). In XPG-deficient cells, (6–4) PPs and CPDs were not removed up to 8 h and 48 h repair time, respectively. By contrast, (6–4) PPs were repaired within 2 h and CPDs within 48 h in XPG-WT cells (**Figure 2E, 2F, S2A, and S2B**). Consistent with the results from UV sensitivity measurements, cells expressing XPG-Δ150, XPG-Δ357, or XPG-Δ654 repaired (6–4) PPs and CPDs simiar to XPG-WT cells, while XPG-Δp62 mutant cells showed delayed repair (**Figure 2E, 2F, S2A, and S2B**). Altogether, these results suggest that XPG interacts with p62-PH through multiple domains and the disruption of those interactions only has a moderate effect on NER activity and UV sensitivity.

### Mutations in XPD binding domain of XPG induce defects UV-induced damage repair

We used an equivalent cell-based approach to assess the importance of the interaction of XPG with XPD for the repair of UV-induced DNA damage. We generated cells with mutations in the XPD-interaction region of XPG (**Table 1 and Figure 3A**) and picked clones with comparable XPG expression levels (**Figure 3B**). We then tested their sensitivity to UV irradiation and the repair kinetics of CPDs and (6–4) PPs. Following exposure to increasing doses of UV-C irradiation, cells expressing XPG-K3E, XPG-4A, or XPG-R3E showed no dramatic loss of survival compared to cells expressing XPG-WT (**Figure 3C**), while cells expressing XPG that combined the three sets of mutations (XPG-ΔXPD, containing K3E, 4A, and R3E) showed significant hypersensitivity to UV (5- to 10-fold, **Figure 3C**). The survival rates of XPG-ΔXPD cells were not statistically different from XP-G patient cells at UV-C doses above 2 J/m^2^ (**Figure 3C)**.

**Figure 3.**
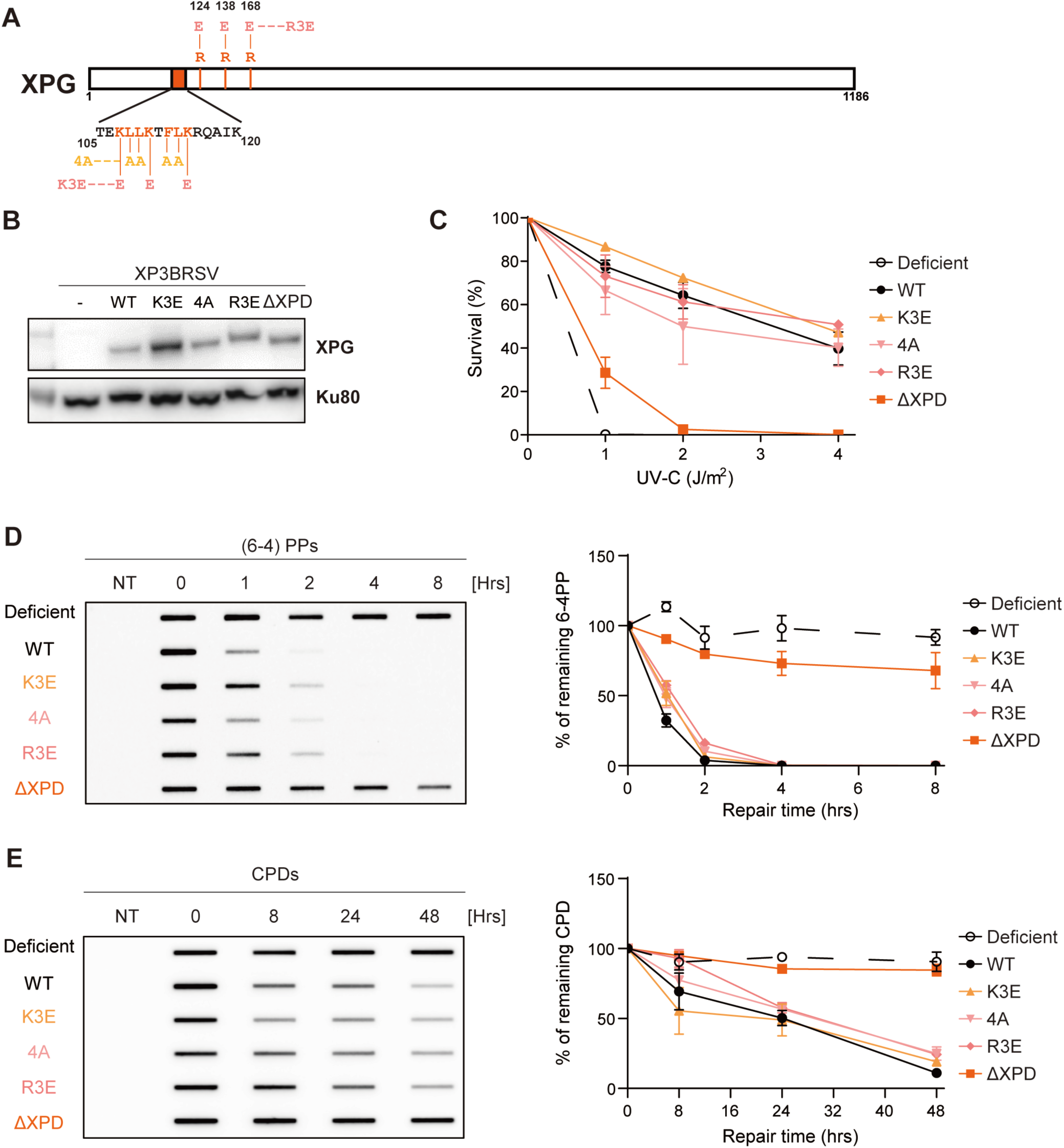
Mutations in XPD binding domain of XPG moderately sensitize the cells to UV. . (A) The XPG domain map with putative XPD interaction regions shown in orange. Predicted XPD interacting basic residues were substituted to glutamic acid, and hydrophobic ones to alanine. (B) Western blot of cells expressing XPD interaction mutants of XPG. Cells were generated by lentiviral transduction and clonal selection to reconstitute mutant XPG proteins in XP3BRSV (XP-G patient) cells. (C) Clonogenic survival assays. Cells were treated with UV-C (0, 1, 2, 4 J/m^2^), grown for 10 days, stained with methylene blue, and colonies counted. Survival rates were normalized to the colony number of non-treated cells and shown as a mean ± SEM of 3 independent experiments. (D) Kinetics of (6–4) PPs repair determined by slot-blot assays. Cells were irradiated with UV-C (5 J/m^2^), genomic DNA isolated at indicated time points, and the adduct levels determined using an anti-(6–4) PPs antibody. For quantification, band intensities were normalized to the WT band at 0 h and shown as a mean ± SEM of 3 independent experiments. (E) Kinetics CPDs repair determined by slot-blot assays. Cells were irradiated with UV-C (5 J/m^2^), genomic DNA isolated at indicated time points and the adduct levels determined using an anti-CPDs antibody. For quantification, band intensities were normalized to the WT band at 0 h and shown as a mean ± SEM of 3 independent experiments.

We next investigated the repair of UV-induced (6–4) PPs and CPDs in the XPG-XPD interaction mutant cells by slot-blot assays. In the XPG-K3E, XPG-4A, and XPG-R3E cells, (6–4) PPs and CPDs were repaired at the same rate as in XPG-WT cells (**Figure 3D, 3E, S3A, and S3B)**. By contrast and consistent with the survival experiments, XPG-ΔXPD expressing cells exhibited a significant defect in the repair of (6–4) PPs, and the level of CPDs repair was indistinguishable from XP-G patient cells (**Figure 3D, 3E, S3A, and S3B)**. In conclusion, our data suggest that XPG interacts with XPD through multiple binding sites and that disruption of the interaction of XPG with XPD sensitizes cells to UV by disrupting NER.

### Interactions of XPG with p62-PH and XPD contribute synergistically to NER

Following our observation that the individual p62- and XPD-interaction mutants of XPG partially abolished NER, we designed XPG variants that combine both mutations to determine if this leads to synergistic effects on the NER activity. We generated cells with mutations in both the p62-PH and XPD interaction interfaces (XPG-Δp62/ΔXPD) and compared their activities with the corresponding single mutants (**Table 1 and Figure 4A**).

**Figure 4.**
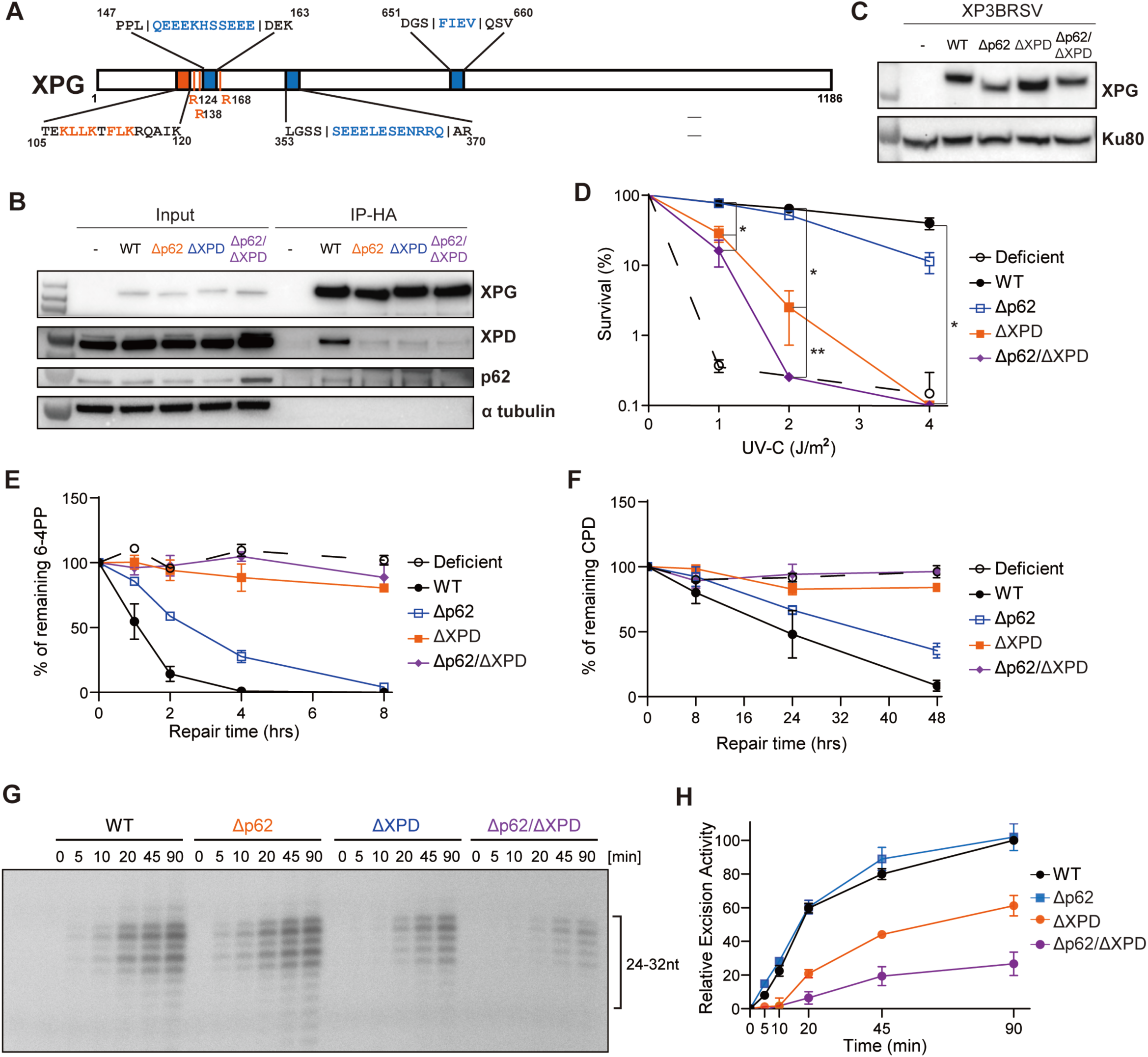
The p62 and XPD interacting domains of XPG synergistically contribute to NER activity. (A) The XPG map with the p62 interaction domains (blue) and the XPD interaction domains (orange) highlighted. (B) Western blot of the input and immunoprecipitation fractions of HA-XPG. XPG, XPD, p62, and α-tubulin antibody were used for detection. A representative figure from 3 independent experiments is shown. (C) Western blot of cells expressing XPG-Δp62, XPG-ΔXPD and XPG-Δp62/ΔXPD generated by lentiviral transduction and clonal selection. (D) Survival rate from clonogenic assays. Cells were treated with 0, 1, 2, 4 J/m^2^ of UV-C, grown for 10 days, stained with methylene blue, and colonies counted. Survival rates were normalized to the number of colonies of non-treated cells and represented by a mean ± SEM of 2 independent experiments. (E) Kinetics of (6–4) PPs repair determined by slot-blot assays. Cells were irradiated with UV-C (5 J/m^2^), genomic DNA isolated at the indicated time points, and adduct levels determined using an anti-(6–4) PPs antibody. For the quantification, band intensities were normalized to the WT band at 0 h and shown as a mean ± SEM of 3 independent experiments. (F) Kinetics of CPDs repair determined slot-blot assays. Cells were irradiated with UV-C (5 J/m^2^), genomic DNA isolated at the indicated time points, and the adduct levels determined using an anti-CPDs antibody. For the quantification, band intensities were normalized to the WT band at 0 h and shown as a mean ± SEM of 3 independent experiments. (G) *In vitro* NER activity of p62 and XPD-interaction mutants of XPG. A plasmid containing a site-specific 1,3-GTG-cisplatin lesion was incubated with purified WT or mutant XPG (27 nM), XPC-RAD23B (5 nM), TFIIH (10 nM), RPA (42 nM), XPA (20 nM), and XPF-ERCC1 (13 nM) proteins for 0-90 min. The excision products were detected by annealing to a complementary oligonucleotide with a 4-dG overhang, which was used as a template for a fill-in reaction with [α-^32^P] dCTP. (H) Quantification of (G). Band intensities were normalized to the product of the XPG-WT reaction at 90 min. The graph is the average product formed from two independent experiments.

First, we tested if our XPG-TFIIH mutations impair TFIIH binding of XPG. HA-tagged XPG was pulled down by HA-conjugated agarose and the TFIIH interaction was assessed using XPD and p62 antibodies. As expected, TFIIH binding of XPG was detected in cells expressing XPG-WT (**Figure 4B**) (58). Mutations in the p62-PH and XPD binding domains clearly reduced TFIIH binding of XPG compared to WT (**Figure 4B**).

We then selected single clones of cell expressing XPG-Δp62/ΔXPD at comparable levels to XPG-Δp62 and XPG-ΔXPD (**Figure 4C**). Cells expressing XPG-Δp62 showed moderate sensitivity to UV and more than 50% of cells survived at 4J/m^2^ of UV-C, while 0% of XPG-ΔXPD cells survived at this dose. XPG-Δp62/ΔXPD double mutant cells showed increased sensitivity compared to single domain mutant cells (**Figure 4D**), showing a 10-fold lower survival than XPG-ΔXPD cells at 2J/m^2^ of UV-C, similar to the XP-G patient cells.

We then measured the repair kinetics of (6–4) PPs and CPDs in XPG-Δp62/ΔXPD cells (**Figure 4E, 4F, S4A, and S4B)**. Consistent to clonogenic survival assay data, XPG-Δp62 showed delayed removal of (6–4) PPs and CPDs, whereas XPG-ΔXPD cells showed residual levels of (6–4) PPs and CPDs repair. Cells expressing XPG-Δp62/ΔXPD were as deficient in (6–4) PPs and CPDs removal as the XP-G patient cells (**Figure 4E, 4F, S4A, and S4B)**. Together, our data suggest that disruption of the interactions between XPG with p62 and XPD synergistically lead to greater NER deficiency compared to individual disruptions.

### Disruption of XPG-TFIIH interactions inhibits NER activity *in vitro*

To assess the intrinsic activity of the TFIIH interaction defective XPG proteins, we analyzed their activity in the *in vitro* NER reactions that are reconstituted with proteins purified from insect cells (**Figure S4C**). We first compared their DNA binding and nuclease activities. DNA binding was measured using a Y-shaped DNA substrate mimicking the ss/dsDNA junction of the NER bubble (**Figure S4D and S4E**). The XPG-Δp62 protein exhibited comparable DNA binding to the XPG-WT protein, while XPG-ΔXPD and XPG-Δp62/ΔXPD showed slightly reduced DNA binding compared XPG-WT (**Figure S4D**). To test the nuclease activity, we used a DNA bubble substrate mimicking the open NER structure. The XPG-Δp62 protein cleaved this substrate with similar efficiency as the XPG-WT protein (**Figure S4E**). Surprisingly, the DNA bubble substrate was not cleaved by XPG-ΔXPD and XPG-Δp62/ΔXPD (**Figure S4E**). We attribute this to the interference with the formation of the “gateway helices”, which are located adjacent to the active site and thought to be important for substrate organization to facilitate cleavage activities (59).

We then measured NER dual incision activity by monitoring the excision of a damage-containing oligonucleotide from a plasmid carrying a site-specific 1,3-GTG cisplatin lesion using the purified core NER proteins (XPC-RAD23B, TFIIH, XPA, RPA, and ERCC1-XPF) in the presence of WT or mutant XPG proteins (**Figure 4G**). The NER activity of XPG-Δp62 was comparable to that of XPG-WT, while XPG-ΔXPD showed a ∼2-fold reduction of the NER activity (**Figure 4G, H**). Incision activity was further reduced with XPG-Δp62/ΔXPD, which displayed only about 20% of the activity of XPG-WT (**Figure 4G, H**).

Intriguingly, the XPG-ΔXPD and XPG-Δp62/ΔXPD proteins displayed significant residual NER activity, despite their defect in nuclease and DNA binding activities (**Figure S4C, S4E, and 4G**). We have previously observed that mutant NER proteins, including the endonuclease ERCC1-XPF, can be inactive on model substrates, yet retain substantial *in vitro* NER activity (22,60). Furthermore, we consistently observe higher activity in the fully reconstituted NER reaction, compared to reactions in cells or cell extracts (61), and has observed the same in the current work. Thus, using the IP assays with TFIIH (**Figure 4B**) and *in vitro* NER assays (**Figure 4G**), our findings indicate that the reduced interaction with TFIIH is responsible for the decreased NER activity observed in the XPG-ΔXPD and XPG-Δp62ΔXPD mutants, which were further substantiated by additional cellular experiments (see below).

### Mutations in TFIIH interaction domain of XPG prevent disassembly of NER preincision complex

Since XPG has both structural and catalytic roles in NER, defects in the interaction between XPG and TFIIH may affect either the association of XPG with TFIIH at the lesion or the formation of a catalytically competent conformation of the complex with XPG. To distinguish these possibilities, we measured the kinetics of the arrival and departure of the XPG variants at sites of local UV DNA damage in cell nuclei. In this experiment, we used CPDs, which are slowly repaired, as a damage marker, and took the residence time of XPG as a measure of the rate of NER observed for (6–4) PPs (33). In XPG-deficient cells, XPG foci were not detected and CPDs persisted 24 h repair time (**Figure 5A and 5B**). In XPG-WT cells, the XPG protein arrived at UV damaged sites within 30 min and was released within 2 h (**Figure 5A and 5C**). XPG was also found at UV-induced damage within 30 min in XPG-Δp62, XPG-ΔXPD and XPG-Δp62ΔXPD expressing cells, indicating that the mutations in the TFIIH interaction domains did not significantly reduce the recruitment of XPG to UV damage. Strikingly, dissociation of XPG from damage was delayed by their reduced interacting with TFIIH. It was completed within 8 h in XPG-Δp62 cells, whereas 20% of XPG-ΔXPD and almost 50% of XPG-Δp62ΔXPD remained bound to CPDs for 24 h (**Figure 5A and 5C**). Overall, our data indicate that the p62-PH and XPD interactions of XPG are required for completion of NER, but not the recruitment of XPG to NER preincision complexes.

**Figure 5.**
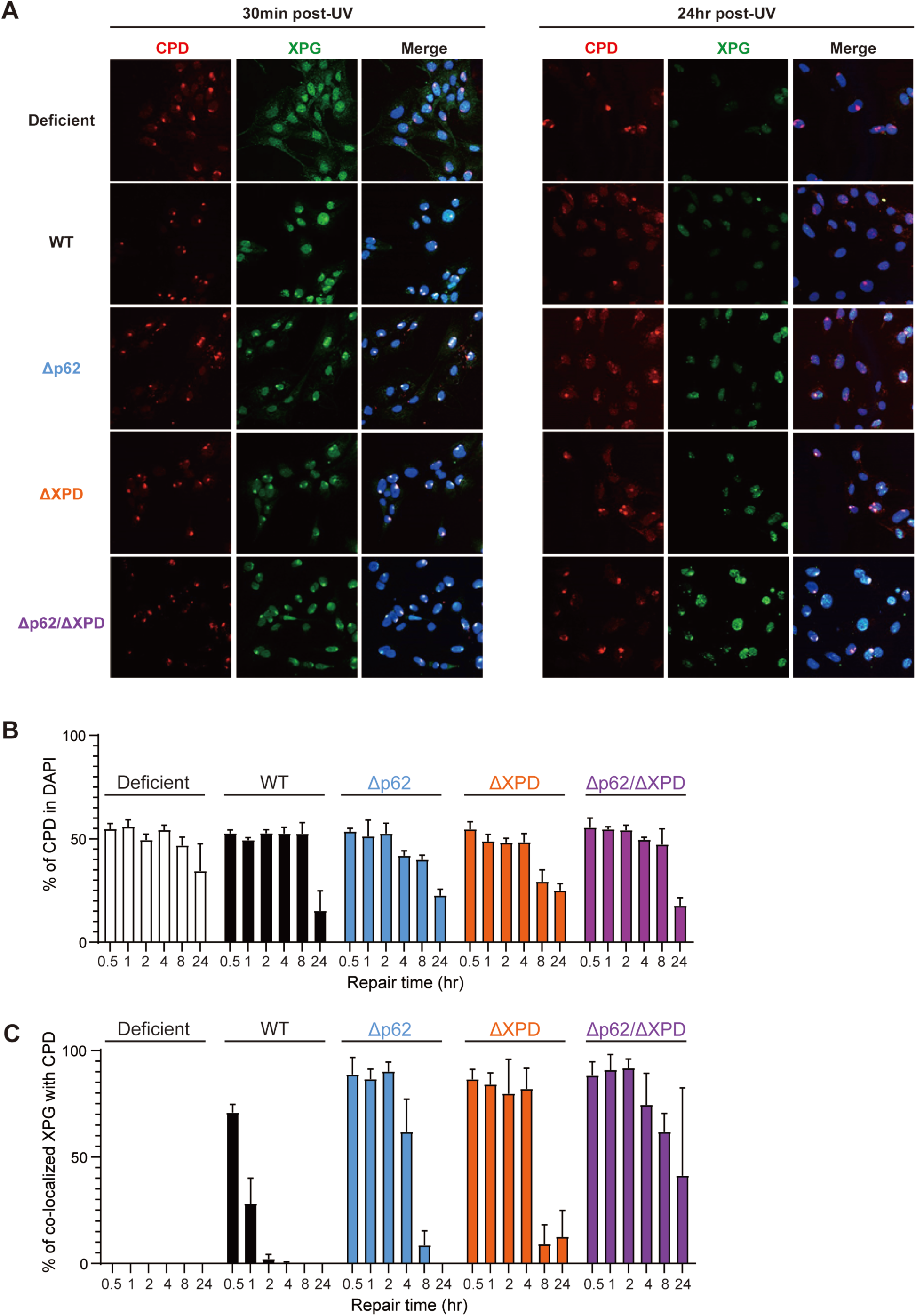
XPG persists at sites of local UV damage in XPG-p62/XPD mutant cells. (A) Representative figure of co-localization assay of XPG and CPDs. Cells were irradiated through a 5 µM micropore filter with UV-C (100 J/m^2^), fixed and stained with HA (XPG, green) or CPD (red) antibodies at different time points. (B) Quantification of fraction of cell nuclei containing CPDs from (A). 100 cells were counted for each sample and data represent a mean ± SEM of 3 independent experiments. % of CPD foci was calculated by dividing number of cells containing CPD foci by the total number of cells. (C) Quantification of XPG co-localization with CPDs from (A). More than 100 cells were counted for each sample and data represent the mean ± SEM of 3 independent experiments. % of co-localization was calculated by dividing number of cells containing CPDs and XPG foci by number of cells containing CPD foci.

### XPD interaction of XPG is needed for incision of XPF

It is known that the presence of XPG, but not its catalytic activity, is required for the 5’ incision by ERCC1-XPF (22,62,63). We therefore investigated whether XPG mutants lacking p62 and XPD interactions affect the dynamics and catalytic activity of XPF in NER. In XPG-deficient cells, XPF localized to UV-induced damage within 30 min and persisted up to 24 h. In contrast, XPF was released from foci in XPG-WT cells within the expected 2-4 h (**Figure 6A, S5A, and S5B**). In XPG-E791A cells, a well-known catalytic mutant of XPG (63), XPF arrived and departed from UV-induced damage as in XPG-deficient cells, indicating that the release of XPF was dependent upon the catalytic activity of XPG (**Figure 6A, S5A, and S5B**). In XPG-Δp62 cells, dissociation of XPF from DNA damage was delayed compared to XPG-WT cells, while XPF remained associated with CPDs until 24 h in XPG-ΔXPD and XPG-Δp62/ΔXPD cells (**Figure 6A, S5A, and S5B**). This suggests that defect in the interaction between XPG and XPD affect the release, but not association, of XPF with sites of UV damage. Lastly, we used COMET Chip assays to ask whether the interaction of XPD and XPG was directly needed to trigger the catalytic activity of XPF to mediate the 5’ incision (51,64). Cells were irradiated with UV-C and single strand breaks (SSBs) were detected by alkaline comet-chip assay (**Figure 6B and 6C**). In the NER reaction, SSBs are transiently formed after incision of the damaged strand and before completion of repair synthesis and ligation. Under normal conditions, gaps are very short-lived and not detected by COMET assays. To render persistent SSBs, cells were pre-treated with DNA repair synthesis inhibitors AraC/HU (65,66). SSBs induction by XPF incision was observed in XPG-WT and XPG-E791A cells as previously reported (64), but breaks were not formed in XPG-deficient cells, formed at intermediated levels in XPG-Δp62 cells, and almost entirely absent in XPG-ΔXPD and XPG-Δp62/ΔXPD cells (**Figure 6B and 6C**). Therefore, the increased time for XPF co-localization with CPDs is correlated with lack of XPF incision activity. This suggests that mutations in the TFIIH interaction domains of XPG are required for proper formation of the preincision complex to trigger XPF incision and that in the absence of XPF incision, NER preincision complexes persist.

**Figure 6.**
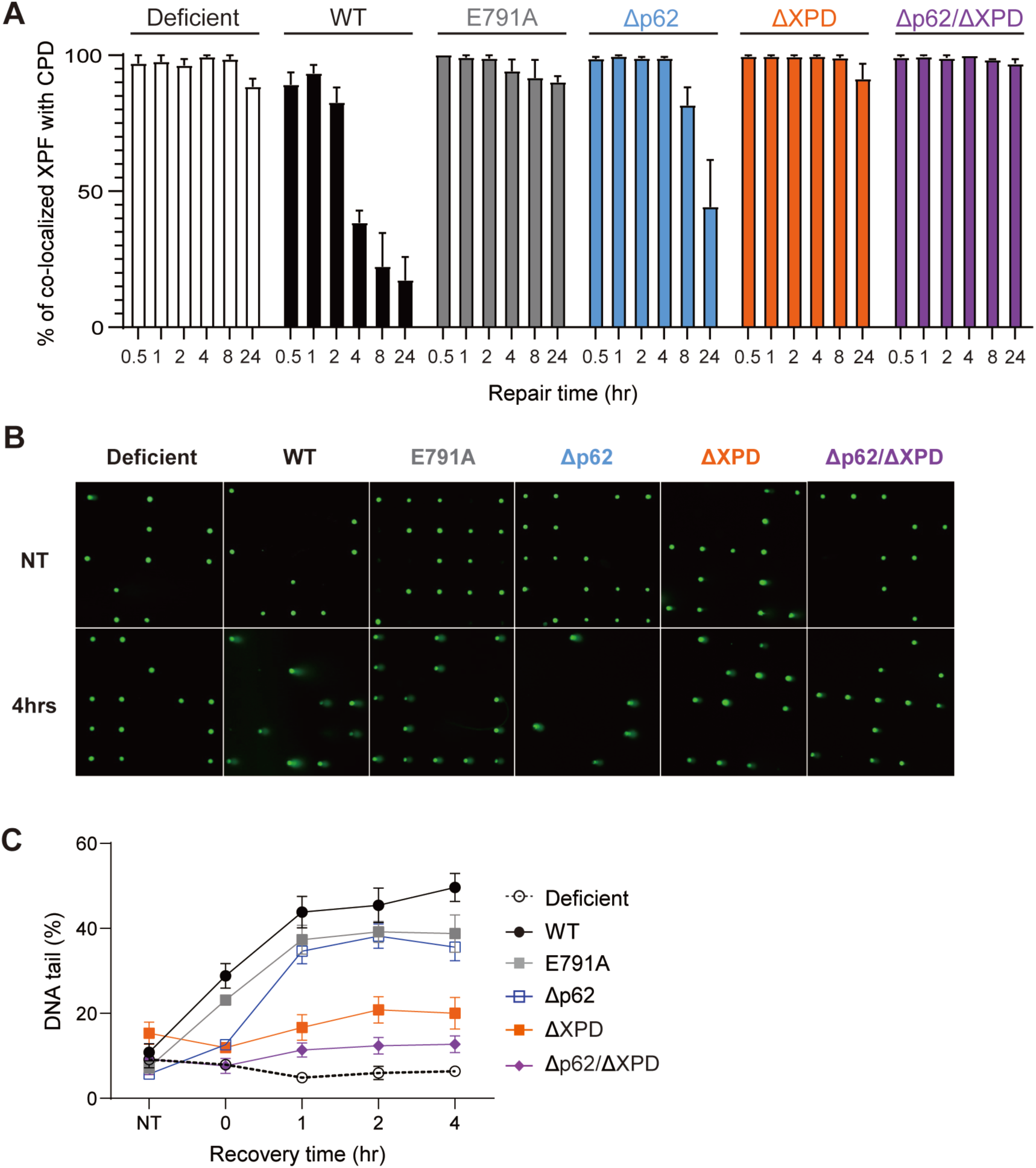
Disruption of the interaction of XPG with TFIIH prevents XPF incision and dissolution of the NER preincision complex. (A) Quantification of XPF co-localization with CPDs. % of co-localization was calculated by dividing number of cells containing CPDs and XPF foci divided by number of cells containing CPD foci. The graph represents the mean ± SEM of more than 3 biological replicates with at least 100 cells each. (B) Representative figure of alkaline comet-chip assays. Cells were pre-incubated with DNA repair synthesis inhibitors (4 mM HU, 40 μM AraC) and irradiated with 5 J/m^2^ of UV-C. Each green dot represents an individual cell and SSBs were represented as tail. (C) Quantification of alkaline comet-chip assays with SSBs quantified as the amount of DNA in the tail on the y axis. The graph represents the mean ± SEM of 3 biological replicates of at least 100 cells each.

## Discussion

Nucleotide excision repair operates through the dynamic and coordinated assembly of the proteins involved at DNA lesions and this process is controlled by protein-protein interactions (25). Among the most complex of these interactions is the one between XPG and TFIIH, owing to a 600 amino acid long unstructured region in XPG called the spacer domain. Although this region is known to interact with TFIIH, it has not yet functionally and structurally characterized. We analyzed the roles of two interaction sites between XPG and TFIIH, specifically with p62 and XPD, in NER at the biochemical and cellular levels. Our data indicates that both interactions are required for full NER activity and contribute to it additively. Disruption of the interaction of XPG with XPD has a more severe effect. Furthermore, we found that these XPG interactions are necessary for proper formation of the preincision complex, which licenses incision by XPF, but not required for recruiting XPG to NER complexes.

One critical transition in NER is the handover from XPC to XPG, as the two proteins occupy overlapping position on the DNA 3’ to the lesion, where XPG ultimately exerts its incision activity (2). Consequently, it has been shown that the two proteins do not coexist in NER complexes (15,67). Since both XPC and XPG bind to the PH domain of the p62 subunit of TFIIH (p62-PH), we hypothesized that this domain may fulfill an important role for this handover and the recruitment of XPG to NER complexes. It is known that an acidic domain in the N-terminus of XPC binds to p62-PH and that this interaction is needed for the recruitment of TFIIH and NER activity (47). Using sequence alignments and taking a cue from yeast studies (44,46), we identified three acidic domains in XPG (amino acids 150-160; 357-368; 651-660) that bind p62-PH with affinities similar to a corresponding peptide of XPC. Contrary to our initial hypothesis, mutations in the p62-interacting domains of XPG resulted only in weak defect in NER, although they acted synergistically with XPD interaction mutants, and did not affect the recruitment of XPG to sites of UV damage. It remains to be determined how these three XPG domains interact with p62-PH in a spatial and temporal manner. Recent computational studies suggested that the acidic patch comprising resides 150-160 may bind to XPD adjacent to an interface of XPD with the “anchor” domain of p62 (56). The full complexities of how the three acidic patches in the spacer region of XPG interact with TFIIH therefore remain to be elucidated.

It is likely that XPG and XPD interact at multiple interaction points, including the recently proposed anchor domain of XPG that may form an extended interaction surface with XPD (56,68). Our analysis here focused on the interaction function for one of the helices in the gateway domain from the active site of XPG (59). The gateway helices are thought to play a crucial role in orienting the DNA at the ss/dsDNA junction for incision (59,69), and have also been modeled to pack against the Arch and FeS domains of XPD (56). We found that multiple mutations in one of the gateway helices (XPG-ΔXPD) severely impaired NER activity. This impairment appears to stem from both a defect in the interaction with XPD and an impact on the catalytic activity (**Fig. 3B and S4E**). Our *in vitro* NER dual incision assays clearly show that XPG-ΔXPD retains partial activity and its activity in NER decreases further with additional mutations in the p62-PH interaction domains (**Fig. 3G, H**). These results indicate that the interaction of XPG with XPD and p62 synergistically contribute to NER.

Interestingly, a reduction in TFIIH binding activity of XPG, caused by mutations in the XPD and p62-PH interaction domains, does not affect the recruitment of TFIIH to NER complexes, but instead, fails to trigger the XPF incision activity and completion of NER (**Fig 6A, C and 7B, C**). In a previous study, the XPG Δ225-231 allele, derived from an XP/CS patient, was defective in TFIIH binding, and this variant could still localize to NER complexes at sites of locally UV-induced DNA damage, albeit with reduced stability at later time points (41). Given that intact TFIIH is required for the proper recruitment (70), additional interaction regions between TFIIH and XPG are necessary for XPG to join the preincision complex (56).

In summary, we show here that the central domain of XPG interacts with the p62 and XPD subunits of TFIIH. This interaction is crucial for completing preincision complex assembly and initiating the 5’ incision by ERCC1-XPF. Our work highlights the modular nature of protein-protein interactions, which make use of multiple interaction surfaces among the involved factors. This modularity allows for a sequential and dynamic progression of NER along the reaction coordinate (25,71)

## Acknowledgements

We thank Areetha D’Souza (Vanderbilt University) for help with the fluorescence anisotropy analysis and insightful discussions. We would also like thank Geunil Yi (UNIST, Korea) for setting up the Alphafold2 station in our lab, Chi-Lin Tsai and John Tainer (MD Anderson, United States) for providing purified TFIIH protein, and Walter Chazin (Vanderbilt University, United States) for purified RPA protein. The fluorescence spectrophotometry was carried out in the UNIST Central Research Facilities (UCRF). The graphical abstract was created with a licensed version BioRender.com.

