## Supplementary Materials for "The interaction of XPG with TFIIH through p62 and XPD is required for the completion of nucleotide excision repair"

**A**

```
S.cerevisiae NEWDLPDIPGFKYDKEDARVNSNKTFEKLMNSING     174
Xenopus      ALYILPPLEDNENNS--SEEEEEREWEERMNQKQR     178
Gallus       AMYVLPSIEDDEKNS--SEEEEEKEWEIRMTQKKL     176
Mouse        TIYVLPPLPEEEKHS--SEEEDEKQWQARMDQKQA     174
Human        TLYVLPPLQEEEKHS--SEEEDEKEWQERMNQKQA     174


S.cerevisiae EDEGCDSDECEWEEVELKPK--------NVK-FVE     386
Xenopus      IQEA---MAESWDHEKHEK--------PSVSGCEA     381
Gallus       IQAA---MVESSSEEELGGEDKEQLN--LTRSVME     388
Mouse        IQAA---MLGSSSEDEPESREGRQSKERNSGATAD     379
Human        MQAA---LLGSSSEEELESENRRQARGRNAPAAVD     379


S.cerevisiae ESSNDENKDDDLEVLSEELFEDVPTKSQ-------     678
Xenopus      IVSDEEFVNEKEDSDSDDSFIEVDSEFSTSNSQHV     705
Gallus       PKEEREGASQSEDSDSDGSFIEVDTVISNEDEFST     627
Mouse        PKPMGPMEMESEESESDGSFIEVQSVVSNSELQTE     669
Human        PKAVEPMEIDSEESESDGSFIEVQSVISDEELQAE     669
```

**B**

| Mutations | ddmut_prediction |
|---|---|
| XPG-K3E | -0.46 |
| XPG-4A | -1.37 |
| XPG-R3E | -0.12 |
| XPG-Δ XPD | -2.11 |

**Figure S1. Sequence alignment of XPG and ddMut score of XPG-XPD mutants.**
(A) Sequence alignment of XPG homologs from *S. cerevisiae*, *Xenopus laevis*, *Gallus gallus*, *Mus musculus*, and *Homo sapiens*. The Tfb1 interaction domains of Rad2 are highlighted in cyan and predicted p62 interacting domains of XPG are highlighted in blue. (B) ddMut score for each of the XPD interacting mutation. The score was predicted using 'https://biosig.lab.uq.edu.au/ddmut/submit_prediction_mm'

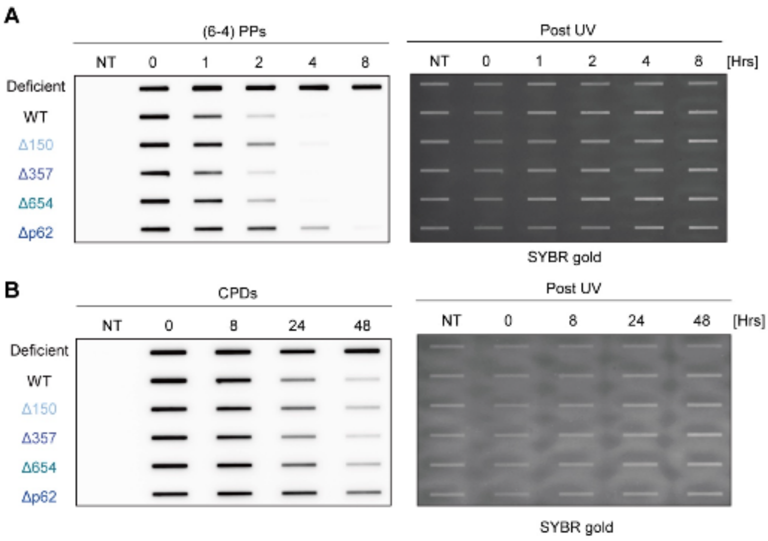


**Figure S2. (6-4) PPs and CPDs repair kinetics of p62 interacting mutants of XPG.**
(A) Kinetics of (6-4) repair determined by slot-blot assays. Cells were irradiated with 5 J/m$^2$, genomic DNA isolated at indicated time points and the adduct levels determined using an anti-(6-4) PPs antibody. Overall DNA levels were determined by staining with SYBR gold to ensure equal loading (B) Kinetics of CPDs repair determined by slot-blot assays. Cells were irradiated with 5 J/m$^2$, genomic DNA isolated at indicated time points and the adduct levels determined using an anti-CPDs antibody. Overall DNA levels were determined by staining with SYBR gold to ensure equal loading.

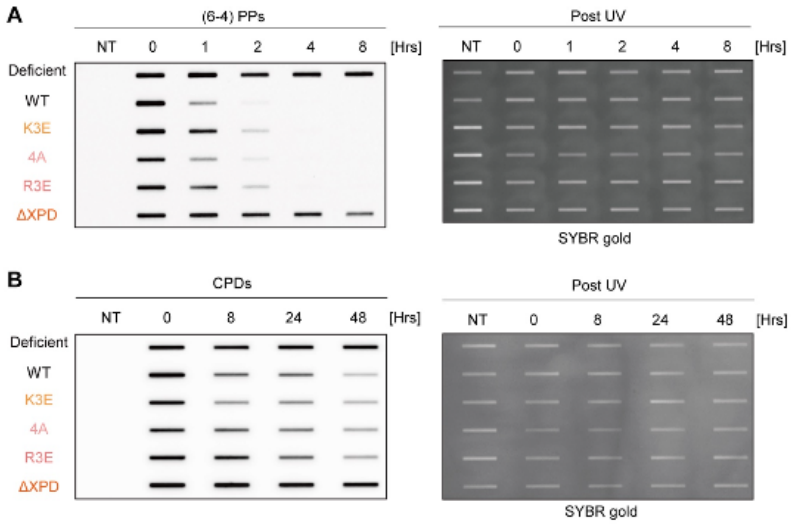


**Figure S3. (6-4) PPs and CPDs repair kinetics of XPD interaction mutants of XPG.**
(A) Kinetics of (6-4) repair determined using slot-blot assays. Cells were irradiated with UV-C (5 J/m$^2$), and genomic DNA isolated at indicated time points and the adduct levels determined using an anti-(6-4) PPs antibody. Overall DNA levels were determined by staining with SYBR gold to ensure equal loading (B) Kinetics of CPDs repair determined by slot-blot assays. Cells were irradiated with UV-C (5 J/m$^2$), genomic DNA isolated at indicated time points and the adduct levels determined using an anti-CPDs antibody. Overall DNA levels were determined by staining with SYBR gold to ensure equal loading.

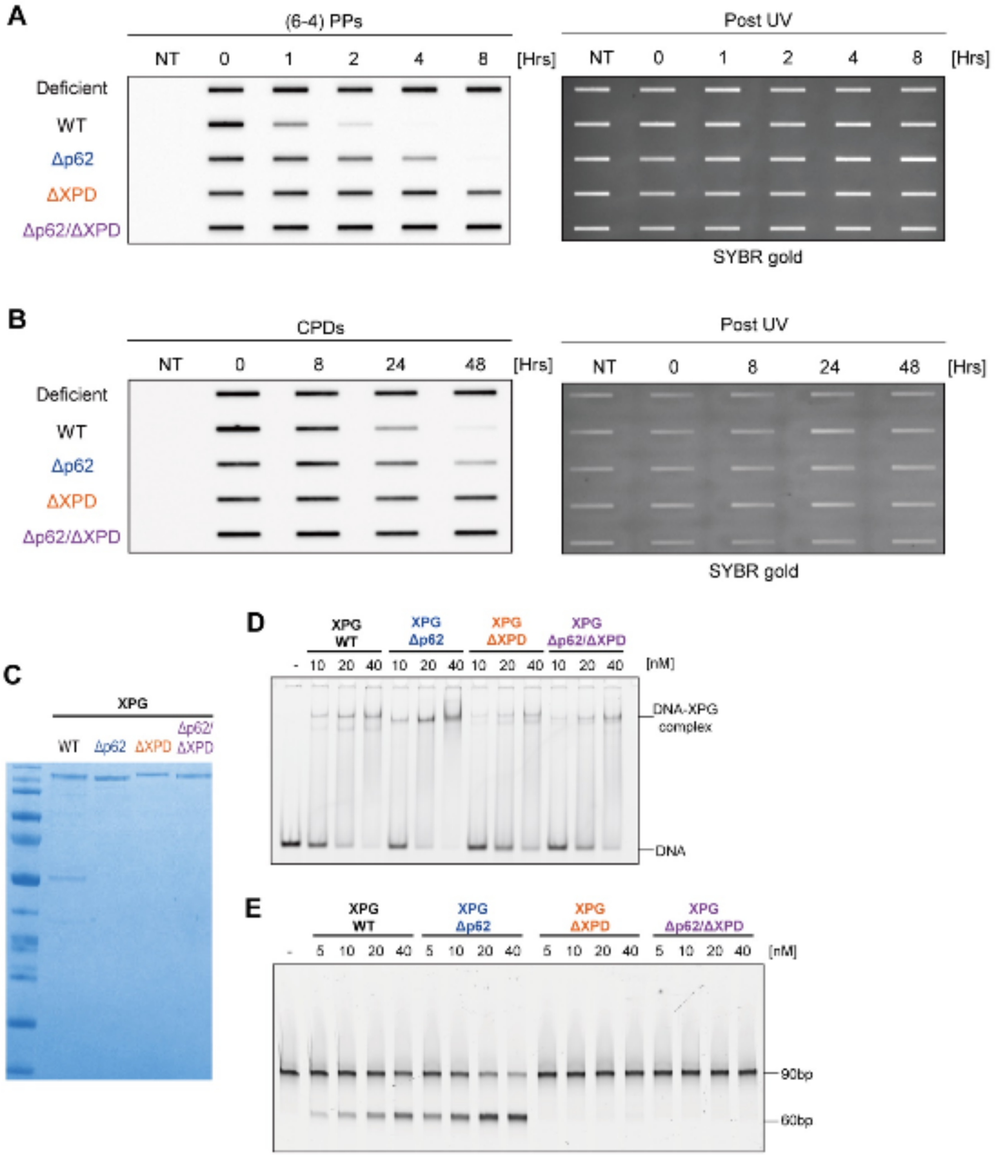


**Figure S4. UV-induced DNA damage repair in XPG-p62/XPD mutant cells and purification of XPG-TFIIH mutant proteins.**

(A) Kinetics of (6-4) repair determined slot-blot assays. Cells were irradiated with UV-C (5 J/m $^{2}$), genomic DNA isolated at indicated time points and the adduct levels determined using an anti-(6-4) PPs antibody. Overall DNA levels were quantified by staining with SYBR gold to ensure equal loading. (B) Kinetics of CPDs repair determined using slot-blot assays. Cells were irradiated with UV-C (5 J/m$^{2}$), genomic DNA isolated at indicated time points and the adduct levels determined using an anti-CPDs antibody. Overall DNA levels were quantified by staining with SYBR gold to ensure equal loading. (C) SDS Page gel showing the purity of the WT and TFIIH-interaction mutant XPG proteins. 0.5 ug of XPG proteins were loaded on the gel. (D) DNA binding assay of WT and mutant XPG with a 5'-FAM-labeled Y-shaped DNA substrate. 5 nM of DNA was incubated with indicated concentration of XPG proteins with 15 nM dsDNA as competitor. (E) Nuclease assays of WT and mutant XPG with a 5'-FAM-labeled 90 mer DNA bubble substrate. 10 nM of substrate was incubated with indicated concentrations of XPG proteins.

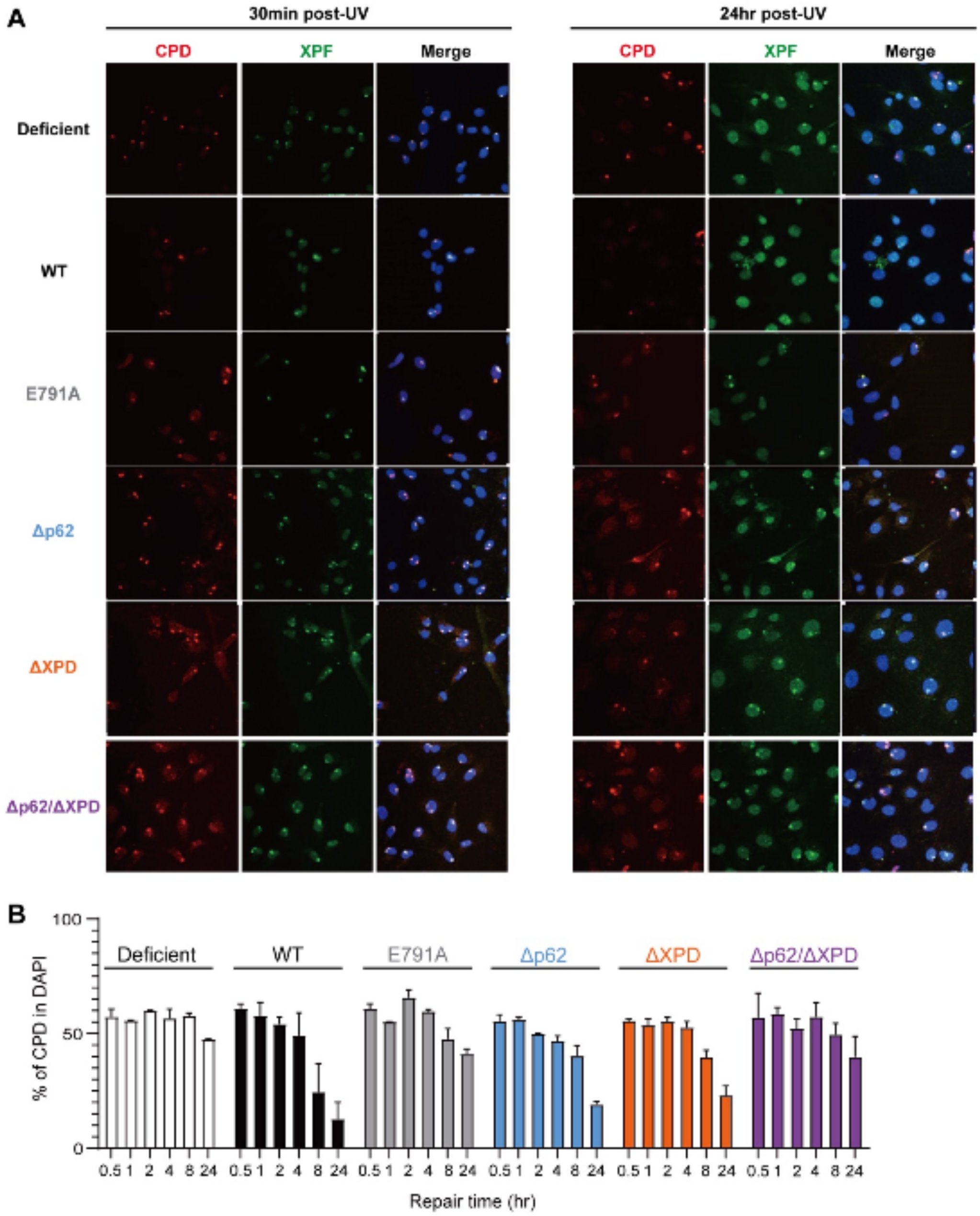


**Figure S5. Disruption of the interaction of XPG with TFIIH prevents XPF incision and dissolution of the NER preincision complex**

(A) Representative figure of co-localization assay of XPF and CPDs. Cells were irradiated through a 5 µM micropore filter with UV-C (100 J/m$^2$), fixed and stained with XPF (green) or CPD (red) antibodies at the indicated time points. (B) Quantification of cells containing CPDs. 100 cells were counted for each sample and data represent 3 independent experiments. % of CPD foci was calculated by dividing number of cells containing CPD foci by number of cells. (Quantification of XPF is in Figure 6A).
